# Murine glomerular transcriptome links endothelial cell-specific molecule-1 deficiency with susceptibility to diabetic nephropathy

**DOI:** 10.1101/143016

**Authors:** Xiaoyi Zheng, Fariborz Soroush, Jin Long, Evan T. Hall, Puneeth K. Adishesha, Sanchita Bhattacharya, Mohammad F. Kiani, Vivek Bhalla

## Abstract

Diabetic nephropathy (DN) is the leading cause of kidney disease; however, there are no early biomarkers and no cure. Thus, there is a large unmet need to predict which individuals will develop nephropathy and to understand the molecular mechanisms which govern this susceptibility. We compared the glomerular transcriptome from mice with distinct susceptibilities to DN, and identified differential regulation of genes that modulate inflammation. From these genes, we identified endothelial cell specific molecule-1 (Esm-1), as a glomerular-enriched determinant of resistance to DN. Glomerular Esm-1 mRNA and protein were lower in DN-susceptible, DBA/2, compared to DN-resistant, C57BL/6, mice. We demonstrated higher Esm-1 secretion from primary glomerular cultures of diabetic mice, and high glucose was sufficient to increase Esm-1 mRNA and protein secretion in both strains of mice. However, induction was significantly attenuated in DN-susceptible mice. Urine Esm-1 was also significantly higher only in DN-resistant mice. Moreover, using intravital microscopy and a biomimetic microfluidic assay, we showed that Esm-1 inhibited rolling and transmigration in a dose-dependent manner. For the first time we have uncovered glomerular-derived Esm-1 as a potential non-invasive biomarker of DN. Esm-1 inversely correlates with disease susceptibility and inhibits leukocyte infiltration, a critical factor in protecting the kidney from DN.

## Introduction

Diabetic nephropathy (DN) is the most common cause of chronic kidney disease and end-stage renal disease in the developed and developing world[1]. Despite clinical trials affirming the importance of glycemic control (in type 1 diabetes) and the inhibition of the renin-angiotensin-aldosterone system (in both type 1 and type 2 diabetes) for slowing progression of nephropathy, beneficial effects are modest[2], and the overall burden of DN continues to increase[3]. Moreover, an increased prevalence of type 2 diabetes[2] and the high cardiovascular risk associated with kidney disease[4] suggest that DN will absorb a disproportionate fraction of health care resources in the coming decades.

Among diabetic complications, DN is associated with the highest cardiovascular morbidity and mortality[5]. However, in patients with either type 1 or type 2 diabetes, the prevalence of nephropathy, defined as macroalbuminuria with or without reduced glomerular filtration rate is less than 15% [6, 7]. Diabetic nephropathy is characterized by changes in the glomerulus including mesangial matrix accumulation[8], podocyte apoptosis[9], endothelial cell injury[10], and leukocyte infiltration[11], all of which can contribute to albuminuria and/or impaired function[8, 11, 12]. While genes responsible for initiation and/or progression of DN have not been definitively identified, genetic predisposition to the disease is thought to play a major role[13, 14].

The NIH-sponsored Diabetic Complications Consortium has phenotypically characterized the clinical response to hyperglycemia among genetically distinct inbred mouse strain[15, 16]. One strain (DBA/2, DN-susceptible) develops albuminuria compared with another more widely used strain (C57BL/6, DN-resistant). Several investigators have shown that DBA/2 mice treated with the β-islet cell toxin, streptozotocin, exhibit increased mesangial matrix accumulation, podocyte apoptosis, and leukocyte infiltration compared with C57BL/6 mice that have similar levels of hyperglycemia[15-17]. The DBA/2 background is also permissive for DN in genetic models of diabetes, Akita and *db*/*db* mice[18, 19].

We hypothesized that by comparing glomerular transcripts from these differentially susceptible mice, we would identify genes that regulate susceptibility to DN. From these transcripts, validation of gene expression and function would inform molecular mechanisms and potential new therapeutic targets for DN.

## Materials and methods

### Animals

We purchased seven week-old DBA/2 and C57BL/6 male mice from Jackson Laboratory and housed these mice in the Stanford University Veterinary Service Center. We then induced diabetes at 8 weeks of age per the Diabetic Complications Consortium protocol [16].with low dose streptozotocin (Sigma Aldrich, St. Louis, MO, USA) for five consecutive days. After four or six weeks, we measured six hour fasting tail vein blood glucose to validate hyperglycemia (Contour, Whippany, NY, USA). We collected overnight mouse urine at the times indicated, and measured urine albumin by ELISA (Exocell, Philadelphia, PA, USA) and urine creatinine by HPLC/MS/MS (Mouse Metabolic Phenotyping Center, Yale University, New Haven, CT, USA). We collected sera by retro orbital bleeding (Kimble Chase, Vineland, NY, USA) immediately before sacrifice.

### Isolation and culture of glomeruli from mouse kidney

We inactivated Dynabeads M-450 (Invitrogen, Carlsbad, CA, USA) with 1 mL 0.1% BSA/0.2M Tris (pH 8.5) at 37°C overnight and then perfused each mouse with 8 × 10^7^ beads in 35-40 mL PBS. We dissociated kidney cells by 1mg/mL collagenase A in 1 mL Dulbecco`s PBS, 37°C for 30 minutes and passed the digested material through a 100 μM cell strainer (Fischer Scientific, Waltham, MA, USA) and isolated glomeruli by a DynaMag-2 magnetic particle concentrator (Invitrogen, Carlsbad, CA, USA). The specificity of isolated glomerular tissue is shown in S1 Fig. We dissolved glomeruli in Tri-reagent (Sigma Aldrich, St. Louis, MO, USA) for RNA preparation, or plated in 1 mL of DMEM (Mediatech, Tewksbury, MA, USA) supplemented with 0.2% FCS (Fischer Scientific, Waltham, MA, USA) in 24-well tissue culture plates and incubated for 24 hours in low (100 mg/dL) or high glucose media (450 mg/dL). We then collected the conditioned media for analysis of Esm-1.

For experiments to separate glomerular and tubulointerstitial fractions, we averted the need for Dynabeads by digesting kidney in 2mg/mL collagenase A in Hepes Ringer Buffer, 37°C for 120 minutes. We then filtered the digested material through a 40 μM cell strainer to collect tubulointerstitial fragments and then filtered the glomeruli on a 100μM cell strainer. We dissolved the glomeruli and tubulointerstitial fractions in Tri-reagent for RNA preparation. The specificity of isolated kidney compartments is shown in **S2 Fig**.

### Microarray analysis

We applied 1 μg of glomerular RNA to an Illumina array (Affymetrix, Santa Clara, CA, USA) to analyze differentially transcribed genes. We compared glomerular RNA from diabetic or non-diabetic DBA/2 or C57BL/6 mice (N=4 / group) and used the non-diabetic, control C57BL/6 mice as the referent group. We submitted the microarray data set to the Gene Expression Omnibus, with accession number GSE84663 (http://www.ncbi.nlm.nih.gov/geo/).

### Gene enrichment analysis

We identified sets of significantly differentially expressed genes, which were mapped to the GeneGo database by MetaCore (lsresearch.thomasonreuters.com). In MetaCore, we calculated the p-value using a hypergeometric distribution. We defined the number of intersecting objects in the experiment as r, the number of network objects in the experiment as n, the total number of intersecting network objects in the database as R, and the total number of network objects in the database as N. We calculated a p-value and false discovery rate for each object in the experiment based on its number of intersections.

### Tissue expression analysis of human esm-1

We obtained publicly available RNAseq data maintained by the European Bioinformatics Institute (http://www.ebi.ac.uk/)[20]. From seven existing datasets, we pooled raw expression counts (Fragments Per Kilobase of transcript per Million mapped reads), pooled similar organs that varied anatomically (e.g. left kidney and right kidney; arm muscle and leg muscle) as duplicates, and compared the median expression across different tissues.

### Quantitative PCR

We used ImpromII Reverse Transcriptase (Promega, Madison, WI, USA) to prepare cDNA from total tissue RNA per the manufacturer’s instructions. We then amplified cDNA by using the qPCR master mix (Applied Biosystems, Grand Island, NY, USA) and the StepOne Plus Real Time qPCR system (Applied Biosystems, Grand Island, NY, USA) with the following protocol: heat activation: 95°C 20s, denaturation 95°C 3s, extension: 60°C 30s, 40 cycles. We used the following primers: Cdh5, forward: TGGTCACCATCAACGTCCTA, reverse: ATTCGGAAGAATTGGCCTCT. CD31, forward: GACCCAGCAACATTCACAGATA, reverse: ACAGAGCACCGAAGTACCATTT. Esm-1, forward: GGCGATAAAACAAGACCAGAAA, reverse: AAACCAGAGATGAGAAGTGATGG. Midkine, forward: AGCCGACTGCAAATACAAGTTT, reverse: GCTTTGGTCTTTGACTTGGTCT. Nephrin, forward: TGCTGCCTTACCAAGTCCAG, reverse: GCTTCTGGGCCGGGTATTTT. SGLT2, forward: TGGCGGTGTCCGTGGCTTGG, reverse: CGGACACTGGAGGTGCCAGATAGC. Tsc22d3, forward: AAGCAACCTCTCTCTTCTTCTCTG, reverse: ATAAGCAGTCATCCCAAAGCTGTA.

### Esm-1 ELISA

The level of mouse serum Esm-1 is approximately 1.0 ng/mL[21]. Therefore, we analyzed mouse serum samples using a Mouse Esm-1 ELISA Kit with a detection range of 23-1500pg/mL (Aviscera Biosciences, Santa Clara, CA, USA). We normalized the urine Esm-1 concentration to creatinine, a quantitative control for glomerular filtration rate. Positive and negative controls for the Esm-1 ELISA are provided in **S3 Fig**.

### Biomimetic microfluidic assay (bMFA)

#### Plating endothelial cells in the microfluidic chip

With the use of our established protocol[22], we coated the chip with fibronectin (100 μg/mL) for 60 minutes and plated human umbilical vein endothelial cells (HUVEC) (Lonza, Walkersville, MD, USA) into the bMFA. After 4 to 6 hours, we applied shear flow to form a three dimensional lumen in the vascular channels. We next activated confluent endothelial cells with 10 units/mL of TNF-α for 4 hours.

#### Leukocyte isolation and labeling

We obtained human blood from healthy adult donors in sodium heparin (BD Biosciences, San Jose, CA, USA), and isolated neutrophils by a one-step Ficoll-Plaque gradient (GE Healthcare, Piscataway, NJ, USA). We next resuspended neutrophils in Hanks Balanced Salt Solution (HBSS) (5 × 10^6^ cells/mL) and labeled with Carboxyfluorescein Diacetate, Succinimidyl Ester probe (Molecular Probes, Carlsbad, CA, USA) for 10 minutes at room temperature. After activation by 10 units/mL of TNF-α for 10 minutes, we incubated neutrophils were with media or recombinant human Esm-1 (R&D systems, Minneapolis, MN, USA) for 10 minutes at room temperature.

#### Leukocyte-endothelial interaction under shear flow

We filled the tissue compartment of the bMFA with chemotactic, N-Formylmethionine-leucyl-phenylalanine (1 μM; Sigma Aldrich) prior to the experiments. The fixed flow rate at 1 μL/min injects 5000 Carboxyfluorescein Diacetate, Succinimidyl Ester probe labeled neutrophils per minute, at 37°C. With a previously developed computational fluid dynamics (CFD)-based model [22], we calculated shear rates in different channels of the network. We recorded video clips at 30 fps using a Rolera Bolt camera (QImaging, Surrey, BC, Canada). After 10 minutes of flowing neutrophils into the bMFA, we injected PBS from the inlet port for 5 minutes to completely wash off unbound neutrophils. We scanned the entire bMFA at the 10× objective using an automated stage on an epifluorescence microscope (Nikon Eclipse TE200, Melville, NY, USA). We processed the acquired images and videos using Nikon Elements software.

#### Data Analysis

We quantified the numbers of rolling, adherent, and migrating leukocytes in the bMFA using Nikon Elements software. We considered any leukocyte traveling at a velocity below the critical velocity as a rolling cell. We estimated critical velocity from a cell velocity flowing in the centerline (v_cc_) as v_crit_ = v_cc_ × ε × (2 – ε), where ε is the cell-to-vessel diameter ratio. We considered any cell that did not move for 30 seconds as adherent.

### Statistics

For analysis of the microarray data, we normalized expression patterns using quantile normalization in R statistical software and deemed differences as significant if there were ≥ 2-fold change between groups if the p-value (adjusted for multiple comparisons using the Benjamini–Hochberg procedure) was below 0.05.

For analysis of human tissue RNAseq data, due to the skewness and heteroscedasity of the raw counts data, we performed the Mann-Whitney test to compare Esm-1, CD31, and Cadherin 5 expression levels in kidney with the other five highest Esm-1-expressing tissues (lung, thyroid, aorta, adrenal gland, and tibial artery). We deemed differences to be statistically significant if the p-value (adjusted for multiple comparisons using a Bonferroni correction) was below 0.05.

For all other experiments, we used Student’s t-test to compare two experimental groups, and one- or two-way analysis of variance to compare three or more experimental groups. We expressed the results as mean +/− standard error of the mean and deemed differences to be statistically significant if the p-value was below 0.05.

### Study approval

The Institutional Animal Care and Use Committee of the Stanford University School of Medicine approved all experiments. For human samples, written informed consent was obtained as approved by the Institutional Review Board of Temple University.

## Results

### Glomerular expression patterns significantly differ between DN-susceptible and DN-resistant mice

To better understand the susceptibility to kidney disease, we compared glomerular gene expression profiles from DN-susceptible vs. DN-resistant mice (DBA/2 vs. C57BL/6 mice, respectively) four weeks after induction of diabetes by low-dose streptozotocin[15, 16]. The classification of these strains as DN-susceptible or DN-resistant was independently validated by blood glucose and urine albumin in separate cohorts (**S4 Fig**).

By microarray analysis of total glomerular RNA, we aligned gene transcripts for each sample by the mean expression in each group in reference to non-diabetic DN-resistant mice. We then focused on gene transcripts in which the mean fold change was greater or less than 2-fold different between diabetic DN-susceptible and DN-resistant mice. This filter narrowed the list from approximately 25,000 to 250 genes. The heat map for this comparison is depicted in Fig 1. To further refine the analysis, we included only genes that were significantly different between diabetic, but not non-diabetic, groups. We identified 22 significantly higher-expressed genes and eight lower-expressed genes between diabetic DN-susceptible and DN-resistant mice (Table 1). Pathway analysis of this set of genes identified the regulation of the immune system as a significant feature of this comparison. Of the processes identified, ranked by lowest p-value, four of the top five were related to inflammation or the immune system (Table 2). In addition to enrichment for immune-related genes, we noted three differentially expressed genes that were not originally classified in the pathway analysis as regulators of the immune system: Esm-1, Tsc22d3, and Midkine[23-25]. This enrichment analysis does not account for directionality. We noted that among these three gene transcripts associated in the literature with inhibition of leukocyte infiltration, expression of Esm-1 and Tsc22d3[23, 24] were significantly lower in DN-susceptible mice. In contrast, a gene transcript associated with promotion of leukocyte infiltration and more severe DN[25], Midkine, was higher in DN-susceptible mice. We validated that these results were consistent between microarray and qPCR experiments (Fig 2, **Table S1**), using distinct primer sites to target the mRNA sequence. To begin to explore these differentially regulated genes with potential significance for DN, we selected Esm-1 for further characterization.

**Table 1.**
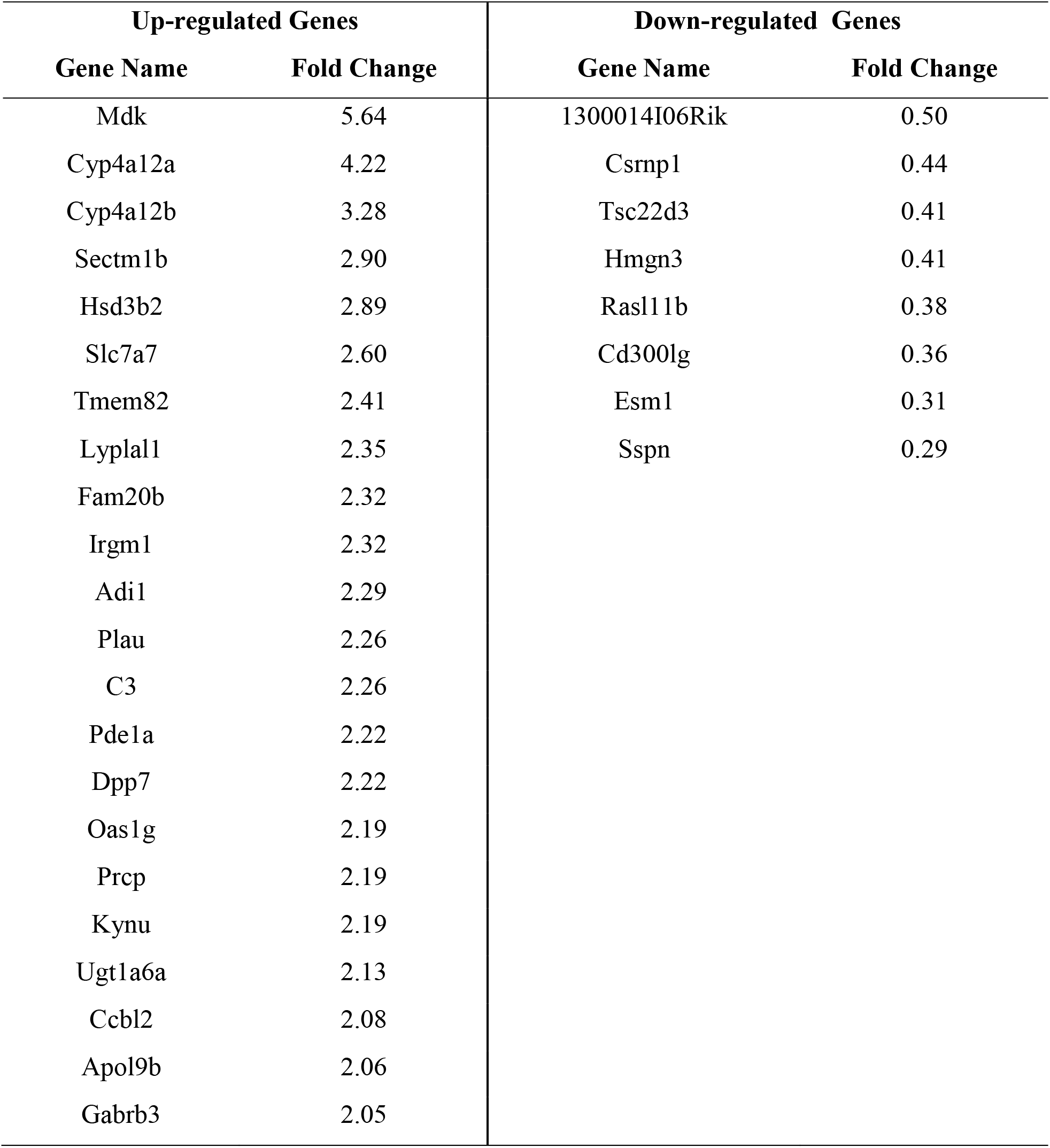
Significantly Up- and Down-regulated genes in diabetic DN-susceptible mice compared to DN-resistant mice.

**Table 2.** Pathway analysis of Up- and Down-regulated genes from microarray.

| Networks | P value | Min FDR* |
| --- | --- | --- |
| Inflammation_complement system | 4.16E-07 | 1.17E-05 |
| Immune response_phagocytosis | 1.23E-03 | 1.72E-02 |
| Proteolysis_Connective tissue degradation | 2.16E-03 | 2.01E-02 |
| Inflammation_Innate inflammatory response | 6.92E-03 | 4.18E-02 |
| Inflammation_Kallikrein-kinin system | 7.46E-03 | 4.18E-02 |
| Proteolysis_ECM remodeling | 1.51E-02 | 6.25E-02 |
| Immune response_Phagosome in antigen presentation | 1.57E-02 | 6.25E-02 |
| Immune response_Innate immune response to RNA viral infection | 1.75E-01 | 4.22E-01 |
| Inflammation_IL-13 signaling pathway | 1.87E-01 | 4.22E-01 |
| Blood coagulation | 1.91E-01 | 4.22E-01 |
| Inflammation_Protein C signaling | 2.17E-01 | 4.22E-01 |
| Inflammation_IL-6 signaling | 2.36E-01 | 4.22E-01 |
| Chemotaxis | 2.67E-01 | 4.22E-01 |
| Immune response_BCR pathway | 2.67E-01 | 4.22E-01 |

\*, Min FDR, Minimum false discovery rate.

**Fig 1:**
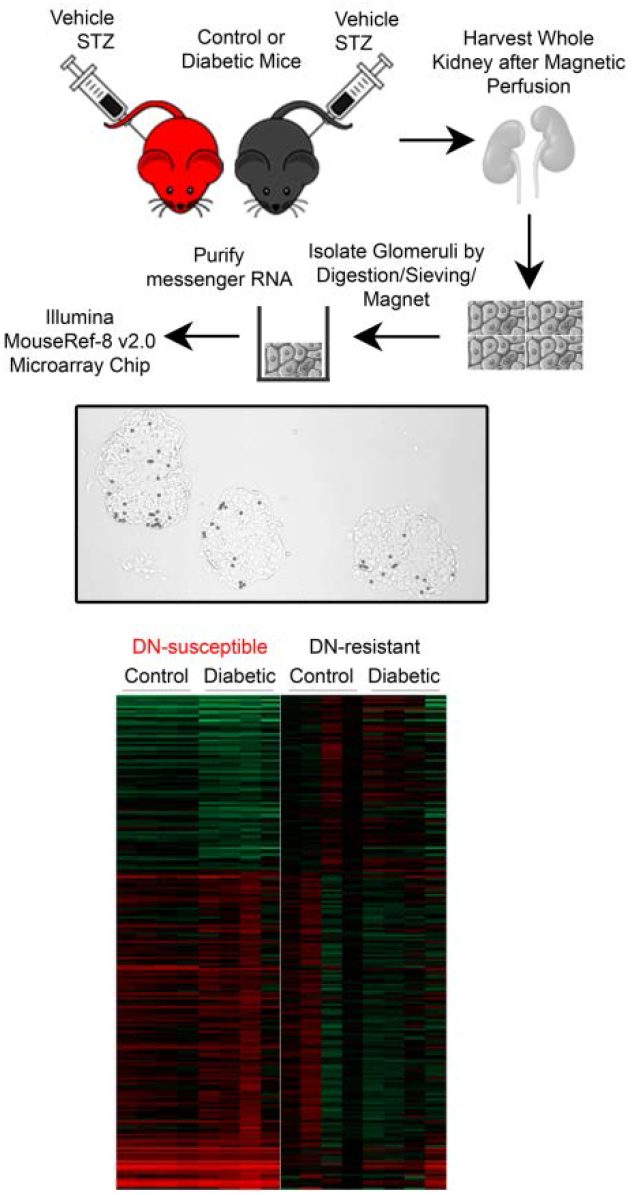
Microarray of differentially expressed glomerular genes between DN-susceptible and DN-resistant mice. (A) Protocol for microarray experiment. DN-susceptible (*red*) and DN-resistant (*black*) mice were injected with vehicle or streptozotocin (STZ), and after four weeks, glomerular RNA was isolated for microarray analysis. (B) Representative brightfield image of isolated glomeruli after magnetic bead perfusion, 200x magnification. (C) A microarray analysis (heat map) aligned by genes for which the mean was > or < 2 – fold more abundant in glomerular RNA from DN-susceptible than DN-resistant mice. Each column represents the values of each sample relative to the mean expression in the control group (glomerular RNA from non-diabetic DN-resistant mice). Each row represents a separate gene from the microarray. Genes with the highest and lowest expression in DN-susceptible vs. DN-resistant samples are shown in *green* and *red,* respectively. Genes colored in *black* were no different from the mean of the control group. N=4 mice per group.

**Fig 2:**
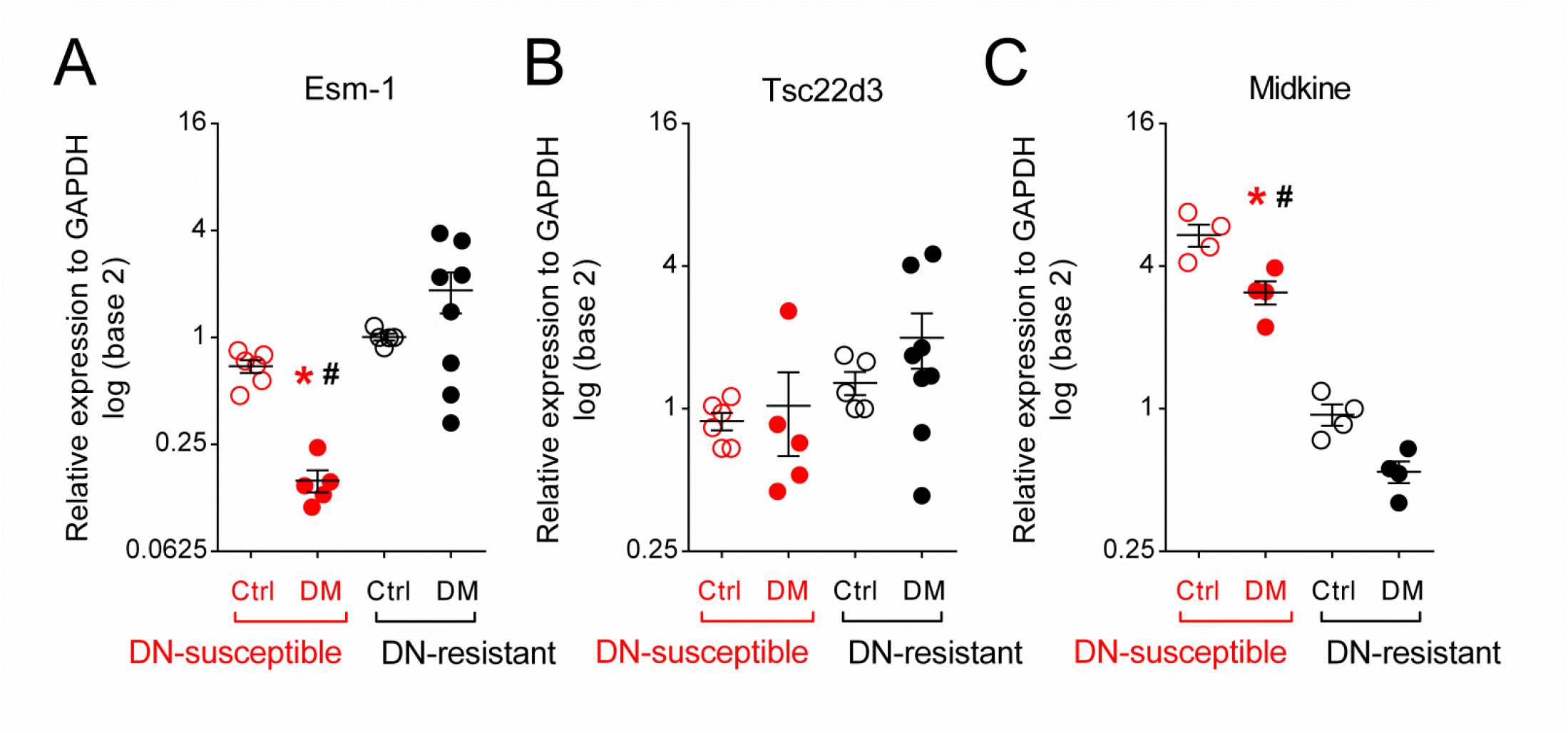
Validating differential expression of 3 immune-related genes from microarray results. Quantitative PCR of relative gene expression from DN-susceptible and DN-resistant mice for: (A) Esm-1, (B) Tsc22d3, and (C) Midkine. Samples from control, DN-resistant mice are used as the reference group. Each circle represents data from one mouse. Open and closed circles indicate data from vehicle- and streptozotocin-injected (i.e. control and diabetic) mice, respectively. Red and black circles/asterisks indicate data from DN-susceptible and DN-resistant mice, respectively. Ctrl, control; DM, diabetic. *, p-value < 0.05 in the same mouse strain between control and diabetic groups. #, p-value < 0.05 in the same treatment between the two mouse strains. N=4-8 mice per group; n=3 replicate wells per sample.

### High expression of Esm-1 and strain-specific difference is found in glomeruli

To ascertain whether Esm-1 was ubiquitously or selectively expressed, we surveyed tissue-specific expression in both humans and mice. We compared Esm-1 expression in datasets from several experiments involving 25 human tissues, and kidney Esm-1 mRNA is the highest, followed by lung (Fig 3A). This difference was not driven by differences in endothelial cell number as two markers for endothelial cells, CD31 and Cadherin 5[26], are not similarly expressed across human tissues. We also compared Esm-1 expression in several tissues from both DN-susceptible and DN-resistant mouse strains, and similar to humans, Esm-1 expression in kidney and lung is highest (Fig 3B). As in humans, this difference was not likely due to a difference in endothelial cell number as CD31 and Cadherin 5 are not similarly expressed across tissues (Fig 3C-D) [26]. Within kidney, Esm-1 expression is significantly enriched for in glomeruli, and is only significantly lower in DN-susceptible vs. resistant mice within this compartment (0.30 ± 0.09 vs. 2.81 ±1.12, p<0.05) (Fig 3E-G). We next examined the regulation of Esm-1 within glomeruli.

**Fig 3:**
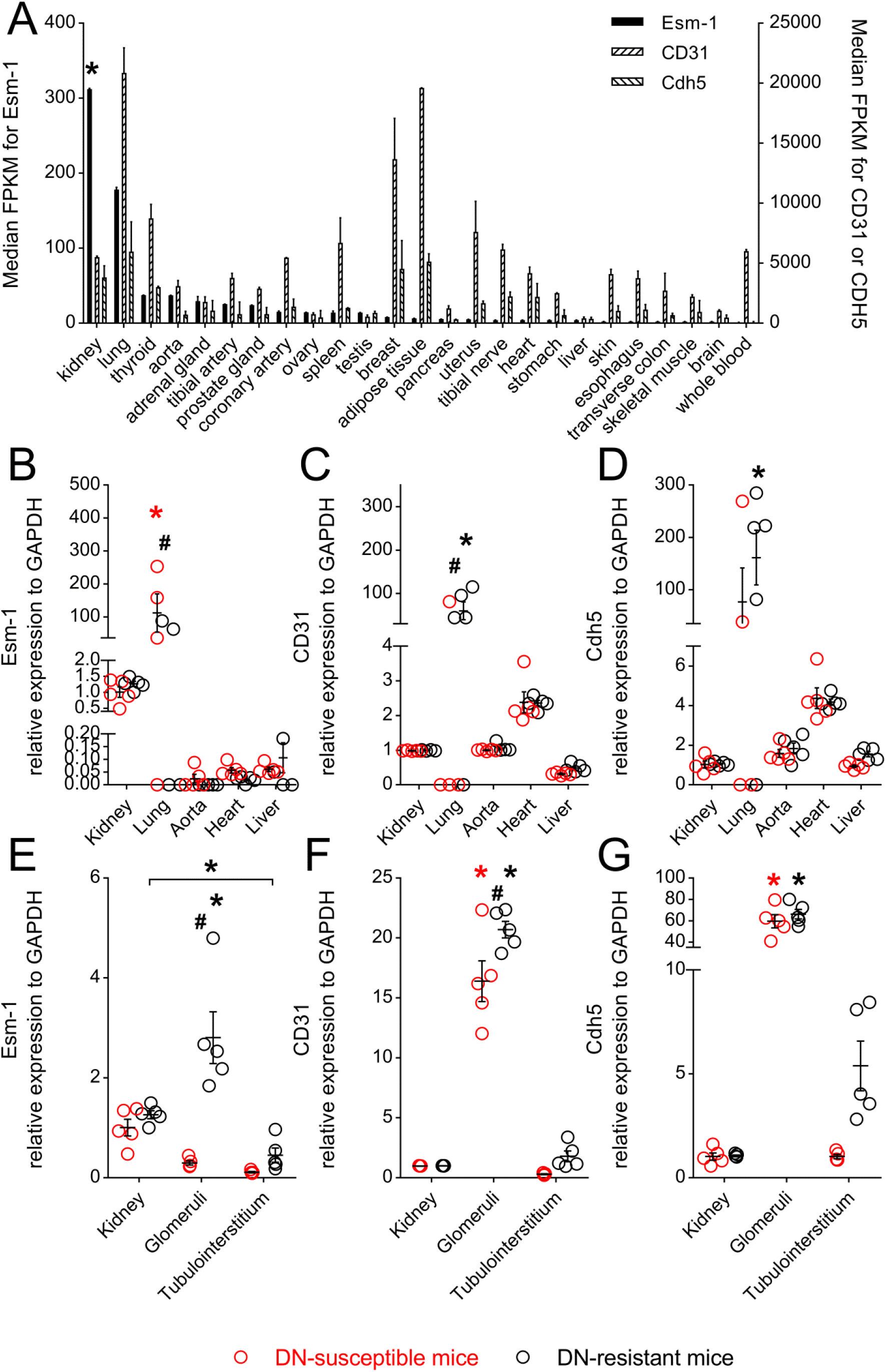
Esm-1 is high in human and mouse kidney and selectively enriched in DN-resistant mice glomeruli. (A) Median Fragments Per Kilobase of transcript per Million mapped reads of Esm-1 (*black*), CD31 (*left hatched*), and Cadherin 5 (Cdh5, *right hatched*) mRNA from seven human tissue RNAseq datasets are shown. Kidney and lung have significantly higher Esm-1 expression than thyroid, aorta, adrenal gland, and tibial artery, and this pattern does not parallel two markers of endothelial cells, CD31 and Cdh5. *, p-value < 0.05 vs. thyroid, aorta, adrenal gland, and tibial artery. (B-G) Quantitative PCR for Esm-1, CD31, and Cdh5 mRNA from select tissues (B-D) and kidney compartments (E-G) is shown. Kidney samples from control, DN-resistant mice were used as the reference group. Each open circle represents data from one mouse. Red and black circles/asterisks indicate data from DN-susceptible and DN-resistant mice, respectively. *, p-value < 0.05 in the same mouse strain between different tissues/compartments; #, p-value < 0.05 in the same tissue/compartment between the two mouse strains. N=4-5 mice per group; n=3 replicate wells per sample.

### Glomerular Esm-1 secretion inversely correlates with DN susceptibility

Esm-1 is predominantly secreted[21], and we were unable to detect Esm-1 in glomerular lysates by Western blot analysis (data not shown). Therefore, to test whether Esm-1 protein expression in glomeruli is regulated by diabetes, we isolated and cultured glomeruli, and assayed for Esm-1 in conditioned media by ELISA. Similar to mRNA expression, four weeks after streptozotocin injection, glomeruli from DN-susceptible mice secreted significantly less Esm-1 than DN-resistant mice (Fig 4A).

**Fig 4:**
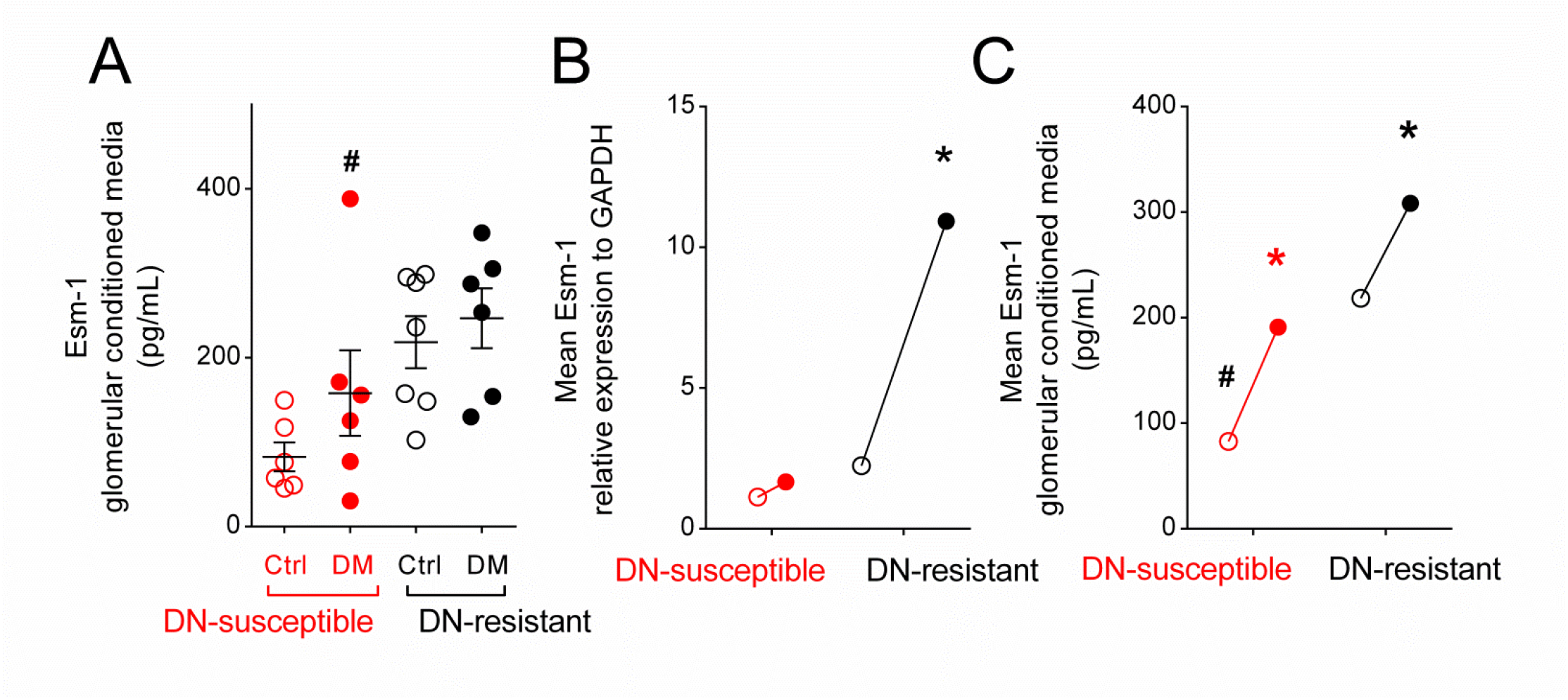
Attenuated secretion of glomerular Esm-1 with diabetes/high glucose in DN-susceptible compared to DN-resistant mice. (A) Glomeruli were isolated from mice treated with vehicle or streptozotocin for 4 weeks and cultured for 24 hours. Mouse Esm-1 was measured by ELISA from conditioned media. Each circle represents data from one mouse. Open and closed circles indicate data from vehicle-injected and streptozotocin-injected (i.e. control and diabetic) mice, respectively. Ctrl, control; DM, diabetic. (B-C) Glomeruli were isolated from mice treated with vehicle only and cultured in low (100 mg/dL) or high glucose (450 mg/dL) for 24 hours. Mean Esm-1 mRNA (B) and secreted protein (C) were compared between low (open circles) and high glucose (closed circles) pairwise from individual mouse glomeruli. Each circle represents the group mean. Red and black circles/asterisks indicate data from DN-susceptible and DN-resistant mice, respectively. *, p-value < 0.05 in the same mouse strain between control and diabetic or between low and high glucose media groups; #, p-value < 0.05 in the same treatment between the two mouse strains. N=6-7 mice per group; n=3 replicate wells per sample in qPCR, and n=2 replicate wells per sample in ELISA.

### High glucose concentration increases glomerular-derived Esm-1

To test whether glucose directly stimulates local glomerular Esm-1 secretion, we isolated glomeruli from non-diabetic DN-susceptible and DN-resistant mice, and assayed for Esm-1 mRNA and protein secretion in low or high glucose media. Incubation in high glucose media increased Esm-1 mRNA and protein secretion in glomeruli from both strains of mice (Fig 4B-C), however the increase was significantly less in DN-susceptible mice.

### Systemic Esm-1 is dynamically regulated in diabetes

The glomerulus is a major source of Esm-1 production (S3 Fig), but the contribution of kidney production to urine or serum Esm-1 has never been tested. We measured urine and serum Esm-1 from control and diabetic mice after 4 weeks of vehicle or streptozotocin injection, respectively. Urine Esm-1 was significantly higher in DN-resistant mice (Fig 5A). To determine the contribution of circulating Esm-1, we measured serum Esm-1 in similar groups of mice (Fig 5B). In contrast to urine levels, diabetes significantly decreased circulating Esm-1. To test the integrity of the glomerular filtration barrier, we measured the urine albumin-to-creatinine ratio, and at 4 weeks after streptozotocin or vehicle injection, the ratio was similar among diabetic and non-diabetic mice from both strains (Fig 5C). Serum creatinine, a surrogate marker for glomerular filtration, was also similar among all groups (Fig 5D).

**Fig 5:**
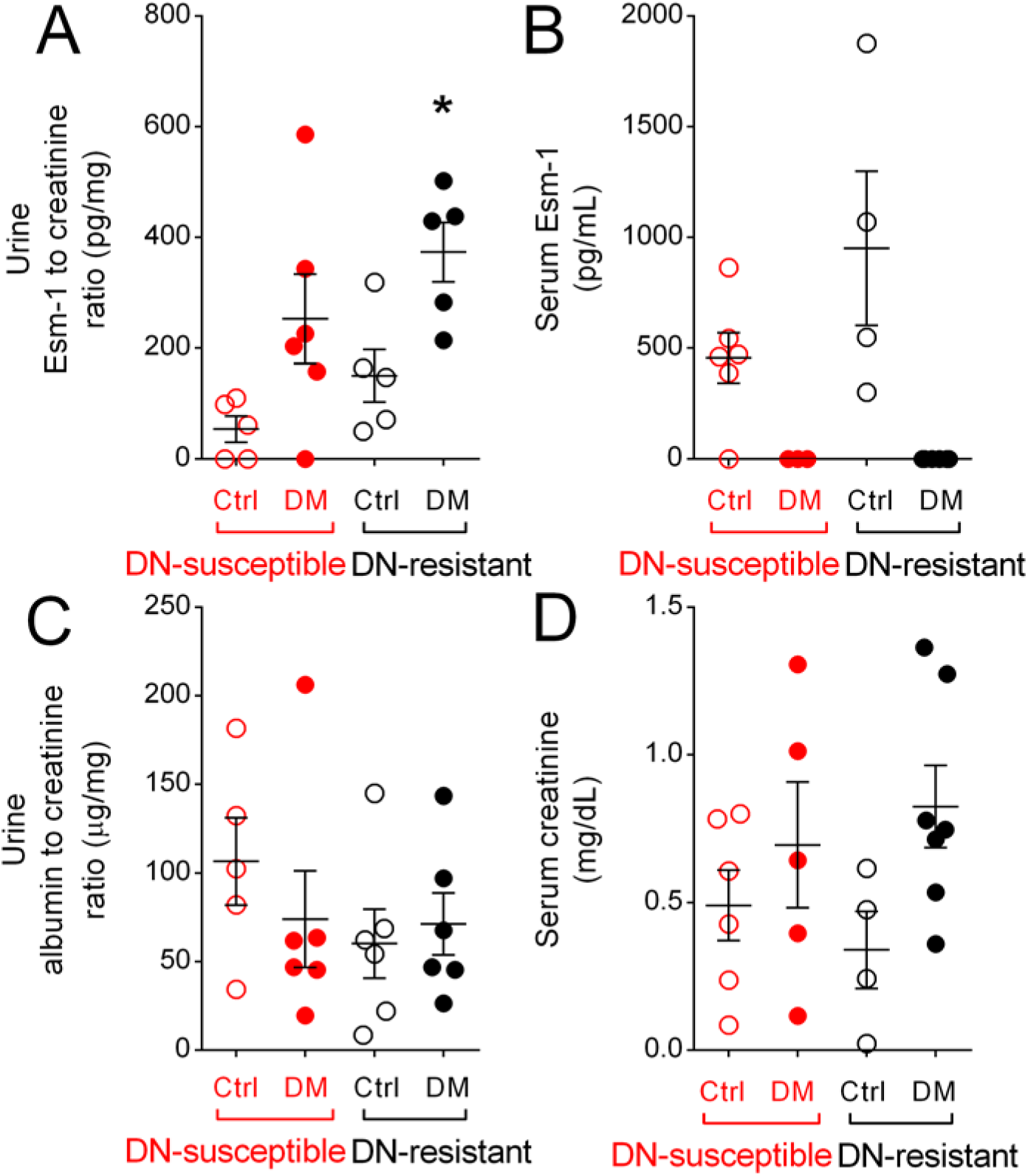
Urine Esm-1 is increased in diabetic mice. Esm-1 level was measured in urine (A) and serum (B) from control and diabetic mice. Urine albumin-to-creatinine ratio (C), a marker of glomerular permeability, and serum creatinine (D), a surrogate for glomerular filtration rate from control and diabetic mice are shown. Each circle represents data from one mouse. Open and closed circles indicate data from vehicle- and streptozotocin-injected (i.e. control and diabetic) mice, respectively. Red and black circles/asterisks indicate data from DN-susceptible and DN-resistant mice, respectively. Ctrl, control; DM, diabetic; *, p-value < 0.05 in the same mice strain between control and diabetic groups; N=3-7 mice per group; n=2 replicate wells per sample.

### Esm-1 inhibits leukocyte transmigration in a dose-dependent manner

To test directly whether Esm-1 blocks leukocyte infiltration across an endothelial monolayer, we utilized intravital microscopy and a biomimetic microfluidic assay (bMFA). Pre-treatment of leukocytes with recombinant Esm-1 showed significantly decreased transmigration (Fig 6A) at 30 and 60 minutes in a dose-dependent manner, suggesting an inhibitory role of Esm-1 against leukocyte infiltration in DN. To investigate the mechanism of decreased transmigration, we examined the role of recombinant Esm-1 to inhibit leukocyte rolling and adhesion (Fig 6B-C). In this *ex vivo* assay Esm-1 did not influence leukocyte adhesion. However, leukocyte rolling was significantly decreased in the presence of recombinant Esm-1 vs. vehicle.

**Fig 6:**
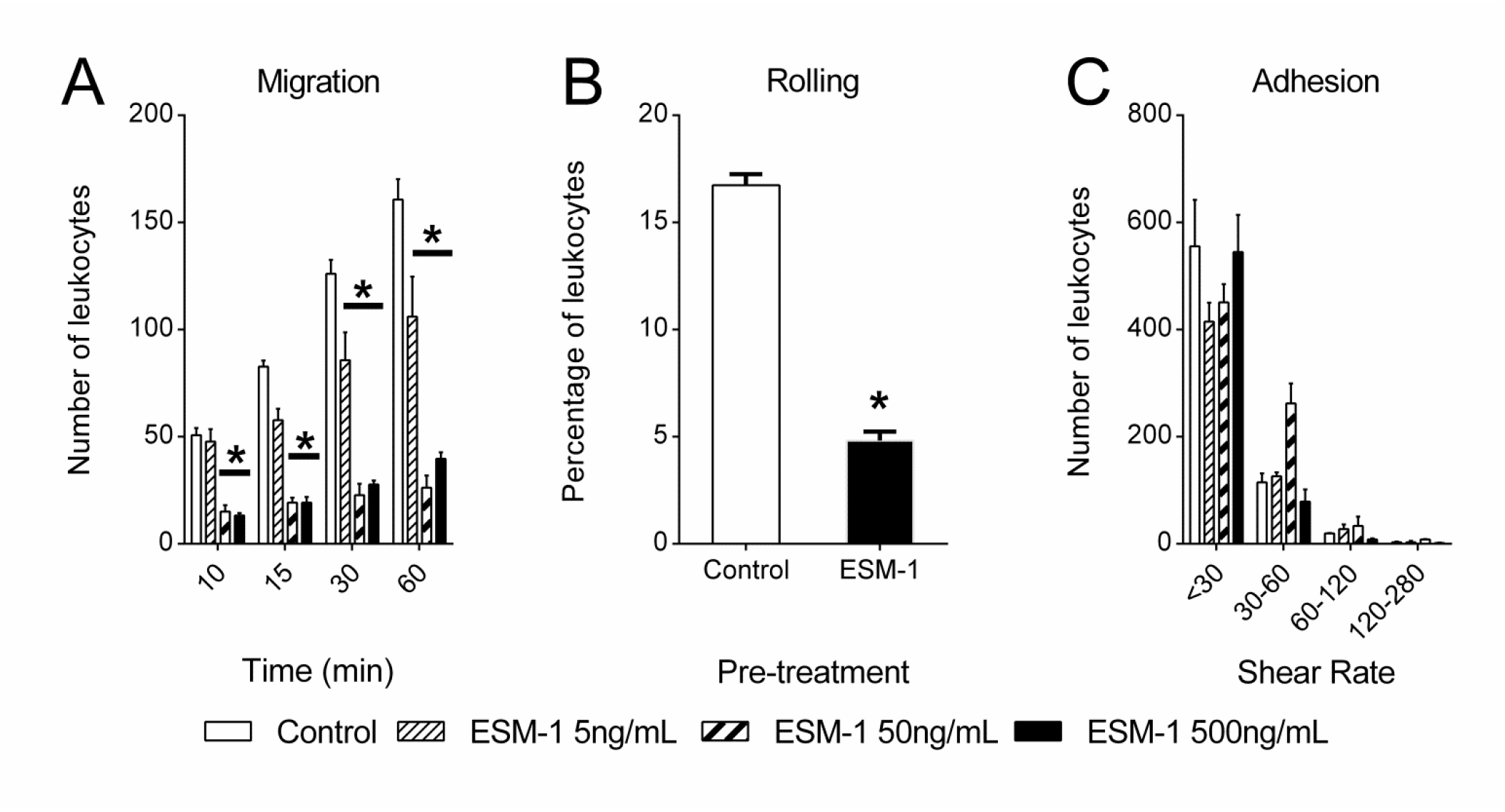
Pre-treatment with Esm-1 inhibits leukocyte infiltration in a dose-dependent manner. A three-dimensional monolayer of endothelial cells is seeded on vessels of a biomimetic microfluidic assay (bMFA) and an activated neutrophil suspension pre-incubated with or without increasing doses of recombinant Esm-1 is injected into the bMFA vascular network. (A) The number of neutrophils transmigrating across endothelial cells at indicated time intervals is quantified relative to untreated neutrophils. (B) The percentage of leukocytes rolling after pre-treatment with vehicle (control) or recombinant Esm-1 is shown. (C) The number of adherent neutrophils on the microfluidic chip is depicted at indicated shear rates. N=3 experiments per dose. *, p-value < 0.05 vs. vehicle group.

## Discussion

We discovered that early expression of glomerular genes related to inflammation is a differentiating marker of susceptibility to DN. Hodgin et al. first demonstrated that differences in glomerular gene transcripts correlate with differences in severity of DN[27]. However, these differences were examined at a late stage of disease, and importantly, compared the response to diabetes in DN-susceptible groups but not the genetic background between susceptible and resistant groups. Based on seminal studies by the Diabetic Complications Consortium [15, 16], we chose DBA/2 and C57BL/6 mice for comparison. Additionally, allele-specific gene sequencing from F1 progeny of these two strains reveals that 41% of genes are differentially expressed in at least one tissue[28]. At an early stage of disease with similar levels of hyperglycemia and when histologic and clinical indices of DN were not yet present[15, 16], DN-susceptible mice have more glomerular leukocyte infiltration compared with DN-resistant mice[17], and our data suggest that leukocyte infiltration is due, in part, to glomerular-specific changes in expression rather than systemic determinants of inflammation.

We sought to identify *glomerular-derived* regulators of leukocyte infiltration as these have not been well characterized. Several lines of evidence suggest that leukocyte infiltration may contribute substantially to glomerular injury, fibrosis, and albuminuria in DN[11, 29-32]. First, glomerular and tubulointerstitial infiltration of macrophages is observed in the diabetic kidney from mice and humans[17, 29, 31, 33], and is proportional to the level of albuminuria[33]. Second, genetic deletion of inflammatory mediators, e.g., leukocyte-attracting chemokine, Monocyte chemotactic protein 1, or intercellular adhesion molecule, ICAM-1[30-32], attenuates the progression of DN in mice. More recently, macrophage-derived factors aggravate glomerular endothelial damage in DN in mice[17]. However, drugs that block leukocyte-endothelial interaction in multiple tissues, e.g., efaluzimab[34], have been removed from the market due to life-threatening infections, highlighting the need for more tissue-specific targeting of the delicate interaction between leukocytes and endothelial cells.

Esm-1 is highly expressed in kidney glomeruli. This relatively high kidney expression was not based on endothelial number, and in fact, the relative difference between mouse kidney and lung could be accounted for by higher levels of endothelial cell markers in lung. However, within mouse kidney, glomeruli had the highest Esm-1 expression likely due to significant enrichment of endothelial cells in this compartment relative to the tubulointerstitium. Interestingly, the strain-specific deficiency of glomerular Esm-1 in DN-susceptible mice was not observed in other tissues. Therefore, we speculate that other glomerular-derived genes may influence Esm-1 transcription rather than a more systemic difference (e.g., Esm-1 promoter sequence) between these genetically distinct strains. Furthermore, glomerular Esm-1 is enriched in endothelial cells[35], consistent with a gene that is native to glomeruli rather than derived from an infiltrating cell. Moreover, lower Esm-1 expression in DN-susceptible mice would be unlikely due to podocytopenia[36]. We further explored the regulation and function of Esm-1 both *in vivo* and *ex vivo.*

Several pro-inflammatory mediators (e.g. TNFα) can induce expression of Esm-1[37], but heretofore, the influence of the diabetic milieu on Esm-1 expression was unknown. Interestingly, this attenuated increase in a susceptible cohort in a *chronic* disease is reminiscent of serum Esm-1 levels in patients with *acute* lung injury, i.e., inflammatory mediators that accompany sepsis can induce Esm-1, but the patients with higher mortality, presumably due to more inflammation, had a smaller increase in Esm-1[38]. The relative deficiency of secreted glomerular Esm-1 in DN-susceptible mice is congruent with the mRNA data but the magnitude of the difference is attenuated. This could be due to alterations in glomerular endothelial cells due to isolation and plating of the glomeruli. We found that after 24 hours, Esm-1 secretion decreased dramatically in culture (**S3 Fig**). Perhaps earlier time points would demonstrate a larger difference between strains of mice, but the detection limit of the ELISA precluded this analysis. Differences in secretion between strains may also be influenced by differences in trafficking of Esm-1 to the plasma membrane or degradation of secreted Esm-1.

To dissect the mechanisms that regulate Esm-1, we demonstrated that high glucose is sufficient to increase glomerular Esm-1 mRNA and protein *in vitro,* and significantly more in DN-resistant compared to DN-susceptible mice. Cytokines that stimulate Esm-1 mRNA (e.g. VEGF) possibly mediate the effect of high glucose as these cytokines are also acutely regulated by high glucose in cultured mesangial cells[39, 40]; however, whether these cytokines participate in DN susceptibility remains an area of further study. Esm-1 transcription is negatively regulated by the transcription factor hHex, but this gene was not differentially expressed in our microarray analysis (data not shown)[41]. Future studies will explore the effects of high glucose on hHex or its binding sites within the Esm-1 promoter and on mechanisms of differential transcription (e.g. promoter methylation) and processing of Esm-1 in glomeruli.

To explore whether this differential expression of kidney Esm-1 is reflected *in vivo,* we measured urine and serum Esm-1. In DN-resistant, compared to DN-susceptible mice, urine Esm-1 was significantly increased with diabetes. Conversely, serum Esm-1 was surprisingly decreased with diabetes. Moreover, a marker of glomerular membrane permeability, urine albumin-to-creatinine ratio, remained unchanged 4 weeks after induction of diabetes. Thus, the filtered load of Esm-1 is not expected to increase. These results suggest that urine Esm-1 may be a candidate non-invasive biomarker of glomerular Esm-1, which increases with hyperglycemia and diabetes and correlates directly with DN resistance. Urine Esm-1 could possibly reflect Esm-1 secretion into the tubule by endothelial cells along the vasa recta. However, tubular secretion would imply transepithelial or paracellular transport of a large ~50kDa protein which is less likely. The mechanism of decreased serum Esm-1 in diabetes is unknown. The diabetic milieu may induce an Esm-1-directed serum protease. The disconnect between urine and serum Esm-1 also suggests that glomeruli may not contribute significantly to circulating Esm-1. Thus, urine Esm-1 is a potential non-invasive biomarker of glomerular Esm-1 production and protection from leukocyte infiltration.

We further characterized the ability of prevent leukocyte recruitment. Esm-1 binds activated leukocyte free antigen-1 (LFA-1), and antagonizes interaction with endothelial-cell expressed ICAM-1[37] in a dose-dependent manner *in* vitro[23]. By using a biomimetic microfluidic assay which includes a vascular network in communication with a tissue compartment to mimic physiological flow conditions[22, 42] we found that recombinant Esm-1 was sufficient to directly inhibit rolling and transmigration *ex vivo.* The lowest, significant dose for inhibition of leukocyte transmigration (5 ng/mL) is within the physiologic range of glomerular-secreted Esm-1 if we assume that the volume of an isolated Bowman’s capsule is 1.5 × 10^5^ μm^3^ [43]. Surprisingly, leukocyte-to-endothelial cell adhesion was not reduced by Esm-1. These data would suggest that LFA-1 is not the only adhesion molecule on leukocytes that binds to ICAM-1[44]. Moreover, these data also underscore the importance of studying leukocyte infiltration in a microfluidic assay, where *in vivo* flow conditions can be modeled[45], and the direct influence on specific mechanisms of leukocyte infiltration (i.e., an effect on rolling, adhesion, and/or migration) can be studied in a vascular network. Our assay was optimized with human neutrophils and human umbilical vascular endothelial cells, rather than macrophages and glomerular endothelial cells, and the role of Esm-1 in leukocyte subtypes and tissue- and species-specific endothelial cells should be confirmed. Moreover, the mechanisms for how Esm-1 decreases rolling remain unexplored.

To our knowledge, this is the first detailed characterization of Esm-1 in diabetes and in kidney with respect to inflammation and DN susceptibility. Esm-1 is enriched in endothelial cells from glomeruli over whole kidney and was higher in glomeruli from *db*/*db* mice vs. controls by IHC[35]. It is conceivable that the *db*/*db* mice used in that study was derived from C57BL/6 vs. KS background (which shares ~14% of the DBA/2, DN-susceptible genome)[19]. Esm-1 was also proportional to markers of inflammation in a cohort of patients with chronic kidney disease[46]. Based on the functional role of Esm-1 that we demonstrated in this study, Esm-1 levels in this prior study may not be an initiator of morbidity but rather a compensatory signal to combat inflammation.

The implications of our work may also extend to acute states of glomerular injury. Glomerular Esm-1 mRNA is decreased in LPS- and anti-GBM-treated mice compared to respective controls[47, 48]. Similar to DN, these acute injury models are ICAM-1 dependent[49], and thus the contribution of a glomerular-derived inhibitor of acute inflammation will be explored in future studies. This strategy is particularly appealing for the pulmonary-renal syndrome of anti-GBM disease[50, 51] as Esm-1 is primarily expressed in kidney but also lung.

In summary, our unbiased screen of glomerular-derived transcripts from DN-susceptible and DN-resistant mice, uncovers Esm-1 as a potential protective gene. This protein is up-regulated in glomeruli and in urine from diabetic *versus* non-diabetic mice, and correlates with resistance to DN. Moreover, *ex vivo* on-chip studies suggest a role for Esm-1 as an inhibitor of leukocyte transmigration across endothelium. These findings motivate further studies of the role of Esm-1 in protecting against DN *in vivo,* and as a marker of resistance to glomerular inflammation in acute and chronic kidney diseases.

## Acknowledgements

X. Z. wrote the manuscript and performed experiments. F. S., E.H., and P.K.A. performed experiments. J.L. performed statistical analysis of the human RNAseq data. S.B. performed statistical analysis of the microarray data. M. K. reviewed and edited manuscript. V.B. conceived the study design and edited the manuscript.

The authors thank Drs. Justin Annes (Stanford University School of Medicine, Department of Endocrinology), Denise Marciano (UT Southwestern Medical Center), Glenn Chertow, and Alan Pao (Stanford University School of Medicine, Division of Nephrology) for helpful discussions.

The authors thank Shripa Patel and Alberto Lovell (Stanford University School of Medicine, Protein and Nucleic Acid Facility) for assistance with qPCR. The authors thank Gary Cline and John Stack (Yale University, Mouse Metabolic Phenotyping Center) for assistance with HPLC/MS/MS.

